# The TCR Cα domain regulates responses to self-pMHCII

**DOI:** 10.1101/2022.05.24.493299

**Authors:** Caleb Y. Kim, Heather L. Parrish, Michael S. Kuhns

## Abstract

T-cells play a central role in adaptive immunity by recognizing peptide-antigens presented in MHC molecules (pMHC) via their clonotypic T-cell receptors (TCRs). αβTCRs are heterodimers, consisting of TCRα and TCRβ subunits that are composed of variable (Vα, Vβ) and constant (Cα, Cβ) domains. While the Vα, Vβ, and Cβ domains adopt typical immunoglobulin (Ig) folds in the extracellular space, the Cα domain lacks a top β sheet and instead has two loosely associated top strands (C and F strands) on its surface. Previous results suggest that this unique Ig-like fold mediates homotypic TCR interactions and influences signaling *in vitro*. To better understand why evolution has selected this unique structure, we asked: what is the fitness cost for development and function of CD4^+^ T cells bearing a mutation in the Cα C-strand? In both TCR retrogenic and transgenic mice we observed increased single positive thymocytes bearing mutant TCRs compared with those expressing wild type TCRs. Furthermore, our analysis of mutant TCR transgenic mice revealed an increase in naive CD4^+^ T cells experiencing strong tonic TCR signals, increased homeostatic survival, and increased recruitment of responders to cognate pMHCII upon immunization, compared to wild type. The mutation did not, however, overtly impact CD4^+^ T cell proliferation or differentiation after immunization. We interpret these data as evidence that the unique Cα domain has evolved to fine-tune TCR signaling, particularly in response to weak interactions with self-pMHCII.

## Introduction

abT-cells play important roles in immunity by scanning the surface of antigen presenting cells (APCs) for peptide-antigens presented by class I or class II MHC molecules (pMHCI/II)(1). CD8^+^ T-cells are restricted to pMHCI and, after activation, differentiate to cytotoxic T lymphocytes (CTLs) that kill infected or transformed cells. CD4^+^ T-cells are pMHCII-restricted and differentiate to a variety of T helper (Th) phenotypes, including Th1, Th2, Th17, or Tfh cells, as well as regulatory T-cells (Tregs), that help direct the responses of other immune cells(2, 3). While it is now apparent that a variety of molecular events contribute to T-cell-mediated immunity, the fine details remain incompletely described.

T-cell recognition of pMHCI/II is mediated by 5-module receptor complexes. The T-cell receptor (TCR) is the receptor module. It directly recognizes the composite surface of peptide-antigens embedded in MHCI/II and relays information about the nature of its interactions with pMHCI/II to associated CD3 signaling models (CD3ge, de, and zz). The CD4 and CD8 co-receptor bind MHCII and MHCI, respectively, in a peptide-independent manner and interact with the Src kinase, Lck, which they sequester away from TCR-CD3 complexes until TCR-CD3 and coreceptor bind and assemble around the same pMHCI/II in a reciprocal manner. The nature of these interactions and the strength of the signals they generate determine if a developing thymocyte undergoes positive selection and commits to the CD4^+^ or CD8^+^ T-cell lineage, or negative selection that induces their death to ensure central tolerance to self-antigens(4). In the periphery, weak tonic signaling to self-pMHC allows naïve T-cells to maintain homeostasis, while strong signals in response to agonist pMHCI/II drive activation and differentiation to effector phenotypes(3, 5). Parameters including the dwell-time of the TCR on pMHCI/II are determinants of the effector phenotype to which they differentiate. In sum, the 5-module complexes that recognize and signal in response to pMHCI/II are central to T-cell development and function.

How pMHCI/II-recognition drives T-cell fate decisions remains an area of active investigation. Considerable effort has been dedicated to determining the architecture of the TCR-CD3 complex, as well as TCR-CD3-pMHCI/II-CD8/4 assemblies, to understand how the spatial relationship of the individual modules impact T-cell responses. Our structure-function analyses, and a recent cryo-EM structure of the TCR-CD3 complex, indicate that CD3ge and CD3de are clustered on one side of the TCR extracellular domain (**Supplemental Figure 1**)(6-8).

Furthermore, contacts between the DE-loop of the TCRa constant region (Ca) and CD3de influence TCR signaling(6). Additional work indicates that CD8 and CD4 are situated near CD3de and CD3ge after co-engagement of pMHCI/II with the TCR, and that the overall architecture of the 5-module assemblies is important for appropriate responses to pMHCI/II(9-11). Finally, because the CD3 heterodimers and coreceptors are situated on one side of the TCR when assembled around pMHCI/II, the Ca surface on the opposite side of the TCR is solvent accessible. Our work has suggested that the C- and F-strands of this surface influence homotypic TCR interactions and responses to pMHCII, leading us to propose that there is a functional sidedness to the TCR(7).

The high conservation of the primary and tertiary structure of this exposed Ca surface also suggests it plays a unique role in T-cell responses. Many activating immune receptors consist of one or more immunoglobulin (Ig) domains wherein two β sheets are held together by a hydrophobic core. Indeed, the variable regions of TCRα and TCRβ (Vα and Vβ) are both Ig-folds, as is the constant regions of TCRβ (Cβ). However, the Cα region of mouse and human TCRs alike consist of a unique Ig-like fold that is missing its top β sheet and instead has two loosely associated top strands, the C and F strands, that form a flexible surface above a hydrophobic core(12, 13). The conservation of this unique Ca domain Ig-like fold in both human and mouse TCRs fold suggests that their solvent exposed surfaces have been selected by evolution because they make an important contribution to T-cell responses.

In this study, we asked: what is the fitness cost of mutating the unusual Cα surface for T-cell development and function *in vivo*? Our goal was to determine how this region normally contributes to T-cell biology. Accordingly, we generated TCR transgenic mice expressing either the wild type (WT) OT-II, or a variant thereof bearing a previously characterized mutation in the Cα domain C-strand. We observed increased positive selection and thymic output in transgenic mice expressing the C-stand mutant when compared to those expressing the WT TCR. Furthermore, we found that naïve T-cells bearing the mutant TCR survive longer than their WT counterparts, and a higher frequency of mutant T-cells proliferated in response to cognate pMHCII. Altogether, our results provide evidence for higher tonic signaling in the mutant CD4^+^ T-cells, suggesting that this unusual Ca surface confers fitness to its host by regulating TCR signaling in response to weak interactions with pMHCII.

## Results

### Mutating the TCR Ca surface alters thymocyte development in retrogenic mice

Before a T-cell can contribute to immunity, its clonotypic TCR must engage self-pMHC well enough to signal positive selection in the thymus, but not so strongly as to signal negative selection. To evaluate the impact of the Cα surface on thymocyte development, we first made retrogenic mice expressing the wild-type (WT) 2B4 TCR (I-E^k^-restricted) or 2B4 mutants bearing the previously described C4 mutation of the C-strand (N151D and K154E) or the F4 mutation of the F-strand (F189E and K196A). Note that residues are numbered by uniprot convention. We also made mice bearing the DE1 mutation of the DE-loop (K171A and D174A), which mediates interactions with CD3δε on the opposite side of the TCR from the unusual Cα surface, as we expected a different phenotype than that of the C- and F-strand mutants. Our analysis showed that, 8 weeks after bone marrow reconstitution, thymocytes expressing the WT and DE1 mutant TCRs were predominantly CD4^+^CD8^+^ double positive (DPs), whereas those bearing the C4 and F4 TCRs were predominantly CD4^+^ single positive (SP). Curiously, the DE1 mutation did not impact thymocyte development compared to WT despite mediating interactions with CD3δε that can impact signaling in response to agonist pMHCII(6, 8). These data suggested that the TCR Cα surface plays a role in thymocyte selection, but did not allow us to evaluate if positive selection, negative selection, or both were being impacted.

### TCR transgenic mice bearing the C4 mutation have an increased SP:DP ratio

To better study thymocytes and peripheral T-cell phenotypes we made TCR transgenic (Tg) mice expressing either the WT OT-II TCR (I-A^b^-restricted), or a variant bearing the C4 mutation (N170D and K173E). These mice were crossed to homozygosity on the C57Bl/6.Rag1 knockout(KO).CD45.1 background. We then generated hemizygous TCR Tg mice on this background, such that one copy would be intact for any endogenous gene or element that was disrupted by the transgene, to mitigate the impact of transgene integration on our analysis. To distinguish our OT-II WT strain generated in the Kuhns Lab from commercially available OT-II TCR Tg mice we will refer to ours as WT^KL^.

First, we asked if the C4^KL^ TCR Tg mice phenocopied the retrogenic mice concerning thymocyte development. The C4^KL^ mutant mice had higher percentage of SP thymocytes than the WT^KL^ even though the WT^KL^ mice had higher numbers of thymocytes overall (**Figure 1B**). These differences were largely attributed to a higher number of DP cells in the WT^KL^ mice compared with the C4^KL^ mice, while the higher frequency of SP thymocytes in the C4^KL^ mice corresponded to a higher absolute number of SP thymocytes compared with the WT^KL^ (**Figure 1C**). The net result was that the C4^KL^ mice had a higher SP:DP thymocyte ratio than the WT^KL^ mice (**Figure 1D**), indicating that these mice did phenocopy the retrogenic mice.

**Figure 1.**
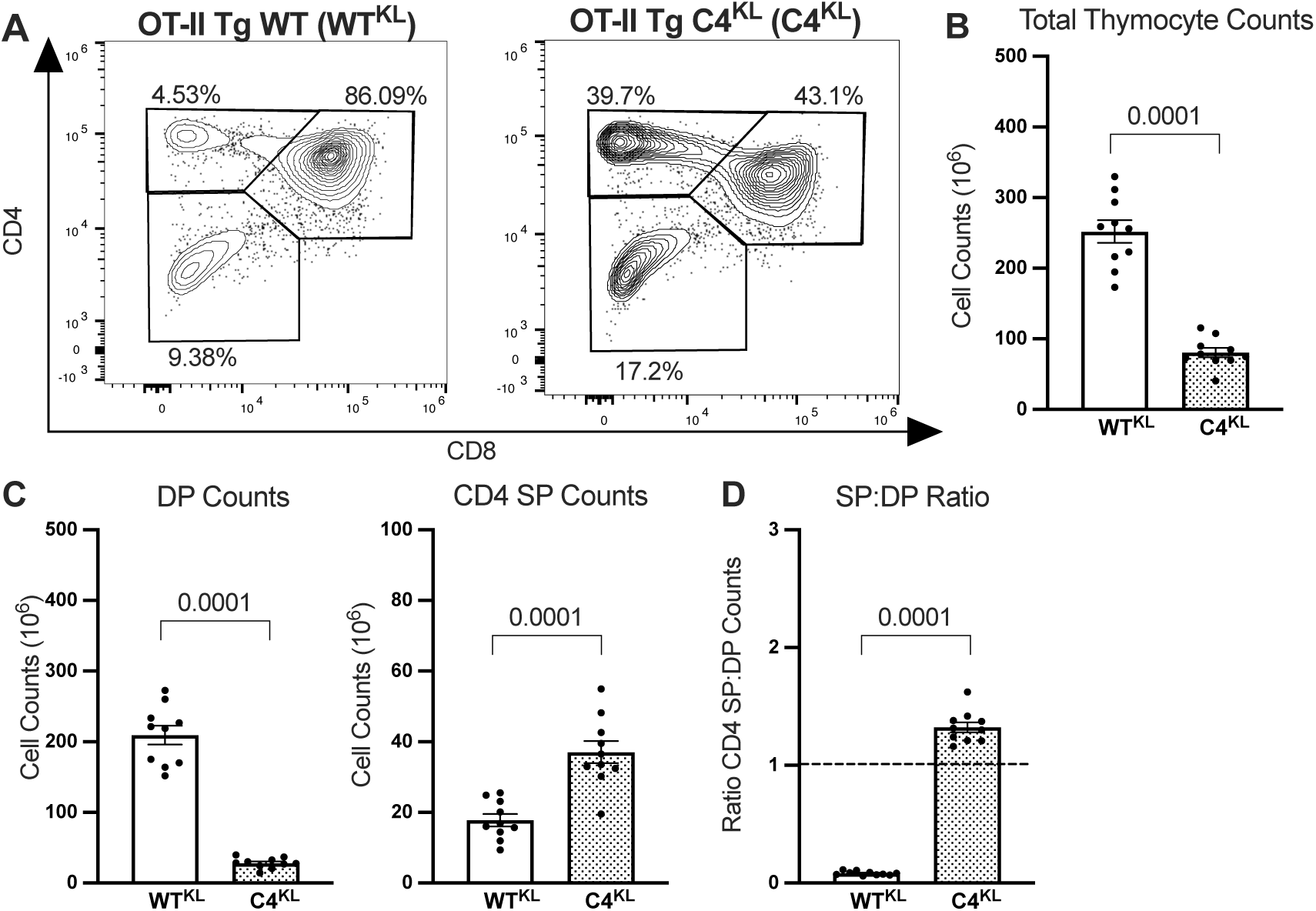
The C4 mutation increases the ratio of SP:DP thymocytes. (**A**) Representative CD8 vs CD4 flow cytometry plots are shown for thymocytes from WT^KL^ (left) and C4^KL^ (right) OT-II TCR Tg mice after lymphocyte gating using SSC vs FSC, FSC-H vs FSC-A doublet discriminator, and Live/Dead Blue (not shown). Percentages are inset for each population. (**B**) The absolute number of thymocytes are shown (right). (**C**) DP (left) and CD4 SP (right) thymocyte counts are shown. (**D**) The ratio of CD4 single positive (SP) counts to CD4^+^ CD8^+^ double positive (DP) counts are shown. Bars represent mean values ± SEM. Each dot represents a single mouse 5-6 weeks of age (male, n=5; female, n=5). Unpaired two-tailed t-test was performed for **B-D** and exact p values are shown.

### The C4 mutant mice have more positively selected thymocytes than WT^KL^ mice

The results introduced above could be explained if the C4 mutant TCR increases the frequency of cells undergoing positive selection relative to the WT TCR. Alternatively, the frequency of thymocytes undergoing positive selection may be equivalent between the mutant and WT^KL^ mice, but the C4 TCR could cause an increased frequency of thymocytes to undergo negative selection. Finally, the results could be explained by an increase in both positive and negative selection.

To investigate the impact of the C4 mutation on positive selection we used gating strategies based on recent publications(14-16). Initially, we gated on CD4^+^ CD5^hi^ thymocytes to analyze both SPs and DPs that had signaled through their TCRs (**Figure 2A**). We then gated on TCR^hi^ CCR7^+^ to restrict our analysis to thymocytes experiencing, or having completed, positive selection. Finally, we used CD69 expression to distinguish thymocytes actively experiencing TCR signaling from those that were not. When we evaluated the CD4^+^ CD5^+^ CCR7^+^ TCR^hi^ CD69^+/-^ populations for CD4 and CD8 expression, we found that the CD4^+^ CD5^+^ CCR7^+^ TCR^hi^ CD69^+^ population were downregulating CD8 but had not fully progressed to the CD4 SP population, suggesting that these cells were auditioning for positive selection(14, 17). In comparison, the CD4^+^ CD5^+^ CCR7^+^ TCR^hi^ CD69^−^ cells had transitioned to the CD4 SP stage, indicating that they had been positively selected (**Figure 2B**).

**Figure 2.**
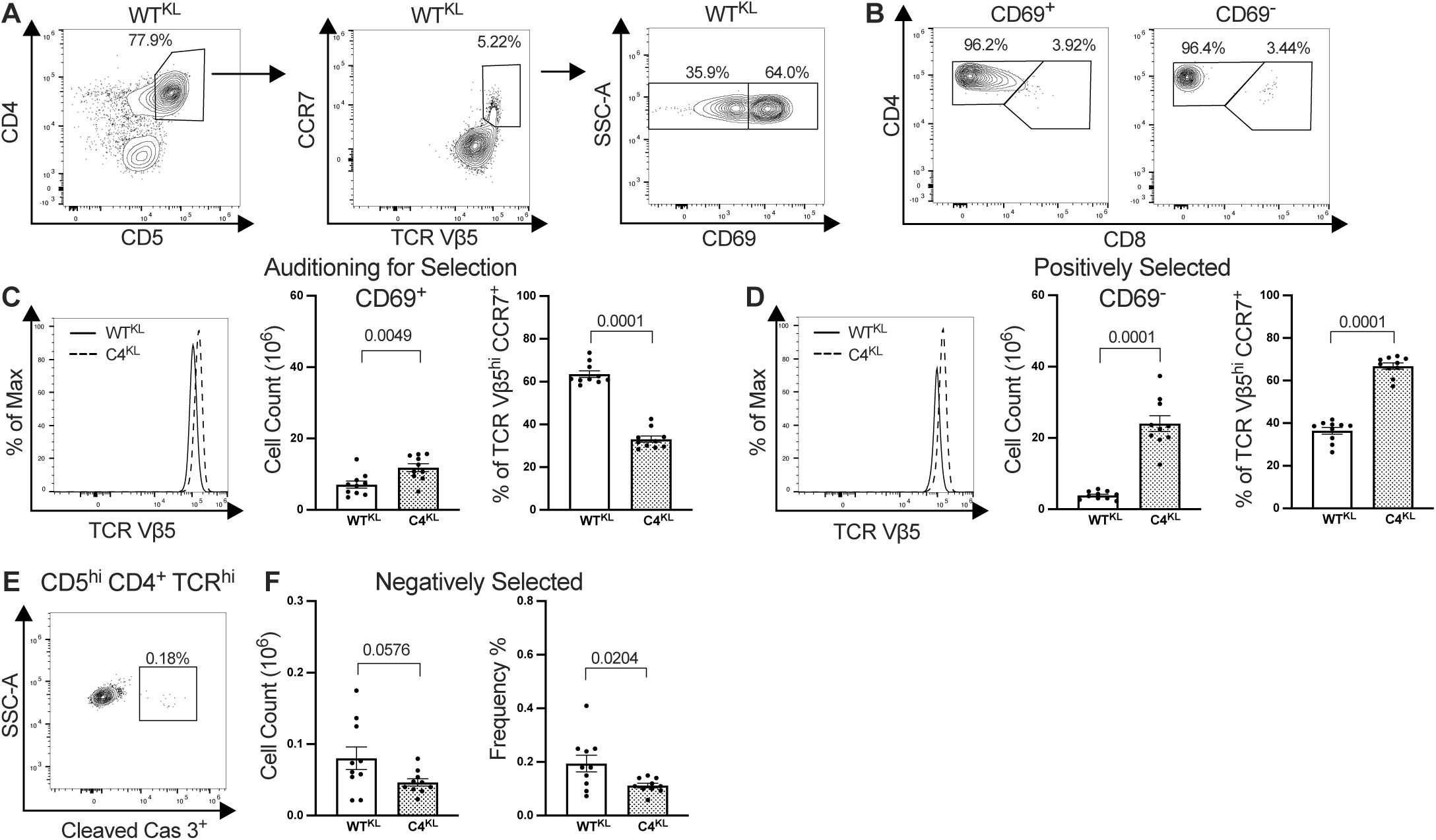
The C4 mutation leads to increased positive selection. (**A**) Representative gating scheme of OT-II Tg thymocytes (WT^KL^ shown as an example), after lymphocyte gating using SSC vs FSC, FSC-H vs FSC-A doublet discriminator, and Live/Dead Blue (not shown). CD4^+^ CD5 ^hi^ WT^KL^ thymocytes (left) were gated based on TCR and CCR7 expression (center) prior to gating based on CD69 expression (right). (**B**) CD8 vs CD4 expression is shown for CD4^+^ CD5 ^hi^ CCR7^+^ TCR^hi^ (Vβ5^hi^) CD69^+^ (left) and CD69^−^ (right) subsets. (**C**) TCR Vβ5 histogram overlays (left), total cell counts (center) and % of parental populations are shown for CD4^+^ CD5 ^hi^ CCR7^+^ TCR^hi^ (Vβ5^hi^) CD69^+^ gated WT^KL^ and C4^KL^ thymocytes subsets. (**D**) TCR Vβ5 histogram overlays (left), total cell counts (center) and % of parental populations are shown for CD4^+^ CD5 ^hi^ CCR7^+^ TCR^hi^ (Vβ5^hi^) CD69^−^ gated WT^KL^ and C4^KL^ thymocytes subsets. (**E**) Representative flow plot for Cleaved Caspase 3^+^ vs SSC-A gating of CD4^+^ CD5^hi^ TCR^hi^ WT^KL^ thymocytes. (**F**) Total cell counts (center) and % of parental populations are shown for of CD4^+^ CD5^hi^ TCR^hi^ Cleaved Caspase 3^+^ thymocytes are shown. Unpaired two-tailed t-test were performed and exact p values are shown. Each dot represents a single mouse 5-6 weeks of age (male, n=5; female, n=5) and bars represent mean ± SEM.

Using this gating we found that, for the CD4^+^ CD5^+^ CCR7^+^ TCR^hi^ CD69^+^ thymocytes, the C4^KL^ mice had overlapping but slightly higher TCR expression compared to the WT (**Figure 2C**). The absolute number of this thymocyte subset auditioning for positive selection was higher in the C4^KL^ mice compared to the WT^KL^ mice; however, as a percentage of the parental CD4^+^ CD5^+^ CCR7^+^ TCR^hi^ CD69^+/-^ population, the C4^KL^ population was lower than the WT^KL^ populations. For the CD4^+^ CD5^+^ CCR7^+^ TCR^hi^ CD69^−^ populations, TCR levels were again overlapping but slightly higher in the C4 mice (**Figure 2D**). Both the absolute number and frequency of this positively selected subset was higher in the C4^KL^ mice than the WT^KL^. Together, these data suggest that a higher frequency of thymocytes undergo positive selection in the C4^KL^ mice compared with the WT^KL^.

To investigate the impact of the C4 mutation on negative selection we analyzed cleaved Caspase3 activity in CD4^+^ CD5^+^ TCR^hi^ thymocytes (**Figure 2E**)(14). We observed reduced numbers of thymocytes undergoing negative selection, which constituted a lower frequency of the CD4^+^ CD5^+^ TCR^hi^ parental population for the C4^KL^ mice compared with the WT^KL^ mice (**Figure 2F**). These data indicate that increased negative selection does not contribute to the increased frequency of CD4 SP thymocytes in the C4^KL^ mice.

### C4 mice have an increased number of recent thymic emigrants and peripheral CD4^+^ T-cells

The enhanced positive selection of thymocytes expressing the C4 mutant TCR described above predicts that there should be increased thymic output, resulting in an increase in recent thymic emigrants. To test this prediction we used a CD24^hi^ Qa-2^lo^ gating scheme to identify recent thymic emigrants (RTE)(**Figure 3A**)(18). Here we added OT-II TCR transgenic mice generated in the Carbone Lab (TCR Tg homozygous, Rag2KO), denoted as WT^CL^, to our analysis(19).

**Figure 3.**
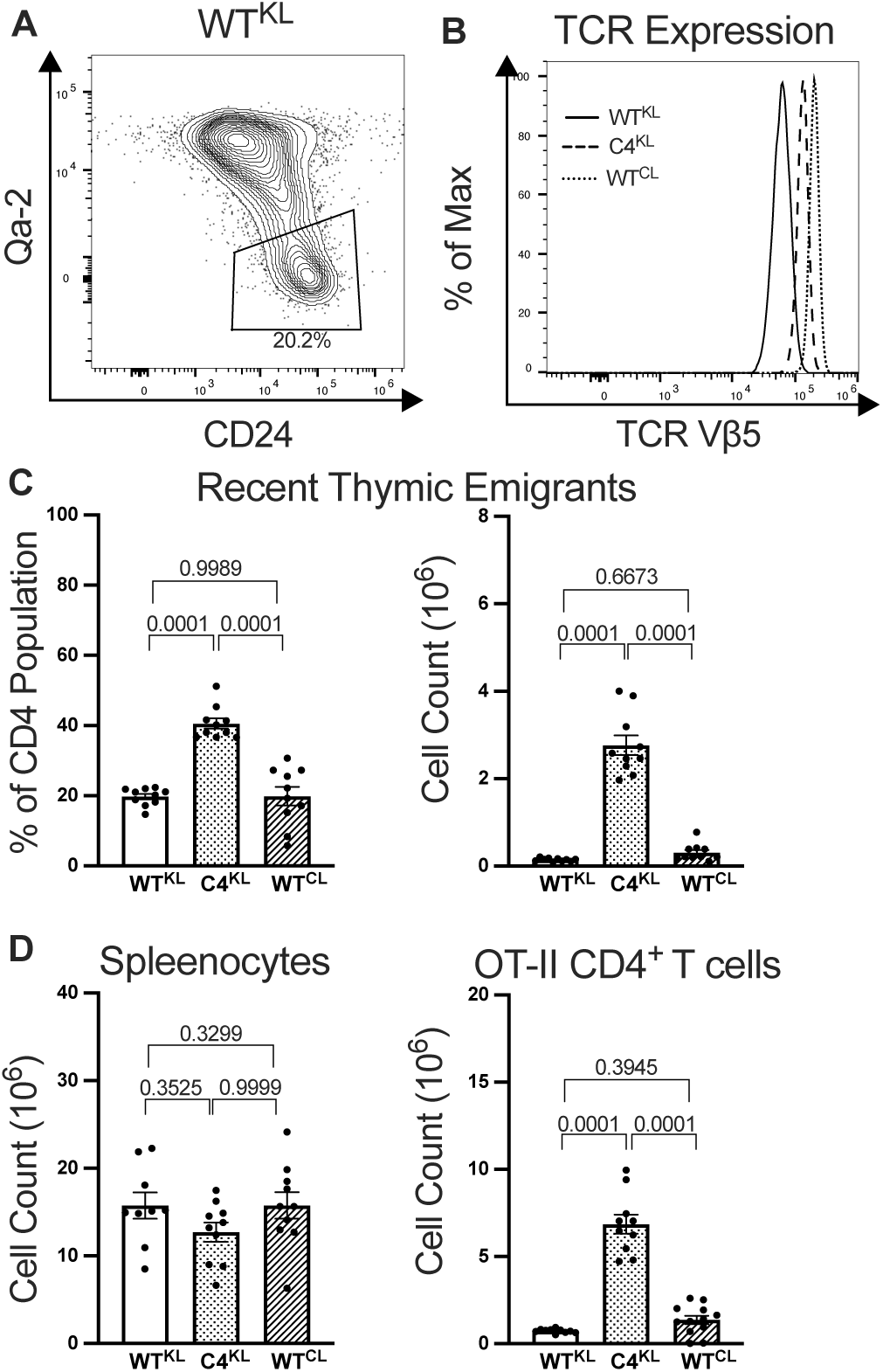
The C4 mutation leads to increased RTEs and peripheral CD4^+^ T-cells. (**A**) CD24 vs Qa-2 expression to show gating for CD24^hi^ Qa-2^lo^ recent thymic emigrants (RTE) after gating on SSC vs FSC, FSC-H vs FSC-A doublet discriminator, Live/Dead Blue, and Vβ5 vs CD4 (not shown). A representative flow plot for WT^KL^ CD4^+^ T-cells is shown. (**B**) TCR Vβ5 levels are shown for WT^KL^ C4^KL^ and WT^CL^ CD4^+^ T-cells. (**C**) The frequency of peripheral CD4^+^ T-cells that are RTEs (left) and their total numbers (right). (**D**) Total spleenocyte counts (left) and CD4^+^ T-cell counts in spleen (right) are shown for WT^KL^ C4^KL^ and WT^CL^ mice. For **C and D**, each dot represents a single mouse 5-6 weeks of age (male, n=5; female, n=5), and bars represent mean ± SEM. One-way ANOVA with Tukey’s post-test were performed and exact p values are shown.

Based on the TCR expression hierarchy on mature CD4^+^ T-cells, WT^CL^>C4^KL^>WT^KL^ (**Figure 3B**), we reasoned that the WT^CL^ strain would help us distinguish between mutation-intrinsic and TCR-expression differences for our analysis of peripheral CD4^+^ T-cell phenotypes and function. We did not include the WT^CL^ mice in our thymocyte analysis because we did not think they were appropriate controls as differences in the transgene constructs, and the timing of transgene expression, could potentially alter thymocyte progression to the DP stages.

Our analysis of CD24^hi^ Qa-2^lo^ RTEs showed that the C4^KL^ mice had ∼2x more RTEs than either strain of WT mice (**Figure 3C**). Interestingly, although we observed an equivalent number of spleenocytes between the three mouse strains, the C4^KL^ mice had nearly 5x the number of CD4^+^ T-cells as the WT^KL^ and WT^CL^ mice (**Figure 3D**). These data are consistent with increased positive selection in the C4^KL^ mice.

### CD4^+^ T-cells in C4 mice experience increased homeostatic survival

The striking increase in mature peripheral CD4^+^ T-cell numbers suggested that, in addition to thymic output, the CD4^+^ T-cells in the C4^KL^ mice may have increased homeostatic survival. Because homeostatic survival is mediated, in part, by weak tonic interactions between the TCR and self-pMHCII, similarly to positive selection, we reasoned that the unusual Cα surface might have evolved to regulate the outcome of these weak interactions.

To explore this further, we first used a CD5 vs. Ly6C gating scheme because CD5^hi^ Ly6C^lo^ CD4^+^ T-cells are reported to experience the highest tonic signaling(20). Our analysis showed that the C4^KL^ mice had a higher percentage of CD5^hi^ Ly6C^lo^ CD4^+^ T-cells compared with the WT^KL^ and WT^CL^ mice, consistent with the idea that the C4 mutation impacts responses to weak TCR-pMHCII interactions (**Figure 4A**). We also asked if we could detect increased basal phosphorylation of the TCR-CD3 complex zeta subunit (pz), as this has also been related to tonic signaling. We did not detect any differences between the strains, however a prior study reported that tonic signaling through the OT-II TCR was so weak that pz could not be detected, which limits our ability to make strong conclusions about this analysis (**Figure 4B**)(21). We therefore tested directly if the C4 mutation leads to increased homeostatic survival by transferring Tag-It Violet-labeled WT^KL^, C4^KL^, and WT^CL^ TCR into C57Bl/6 mice and assessing their survival after two weeks. We recovered 2-10 fold more C4^KL^ CD4^+^ T-cells than WT^KL^ or WT^CL^ CD4^+^ T-cells (**Figure 4C**). Together, the data in Figures 4A and 4C support the idea that the C4 mutation makes CD4^+^ T cells more sensitive to weak TCR-pMHCII interactions, resulting in increased homeostatic survival.

**Figure 4.**
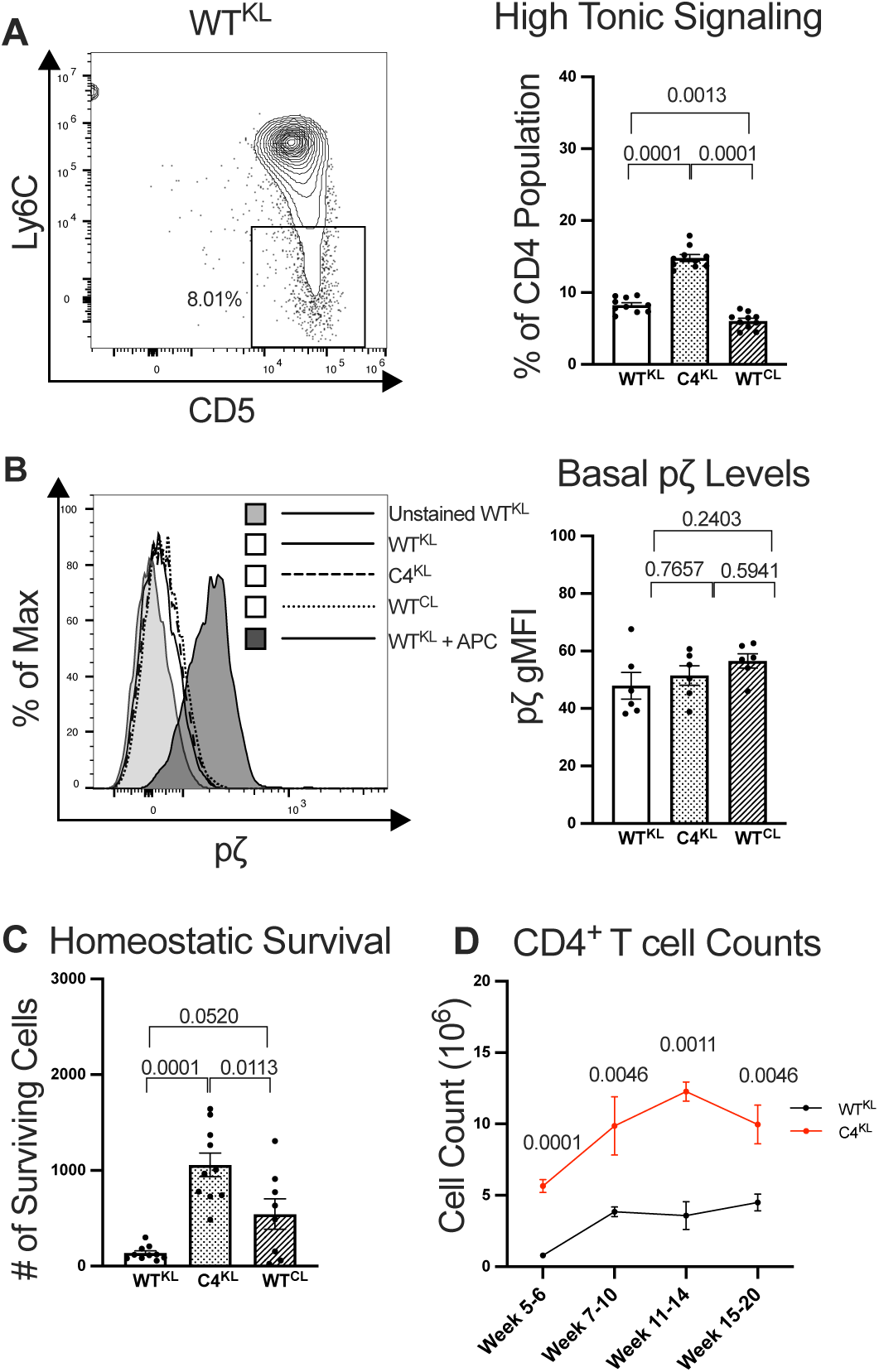
The C4 mutation increases markers of tonic signaling and homeostatic survival. (**A**) CD5 vs Ly6C expression is shown for gating on CD5^hi^ Ly6C^−^ CD4^+^ T-cells experiencing high tonic signaling (left) after gating on SSC vs FSC, FSC-H vs FSC-A doublet discriminator, Live/Dead Blue, and Vβ5 vs CD4 (not shown). Representative WT^KL^ CD4^+^ T-cells are shown. The frequency of CD4^+^ T-cells that are CD5^hi^ Ly6C^−^are shown (right). (**B**) Histogram showing phosphorylation of CD3ζ (left) and gMFI (right) is shown for the indicated populations *ex vivo* or upon stimulation with OVA:I-A^b+^ APCs. (**C**) Quantification of Tag-It violet-labeled WT^KL^, C4^KL^ or WT^CL^ labeled CD4^+^ T-cells 2 weeks after transfer into C57BL/6J recipients. (**D**) Enumeration of CD4^+^ T-cells from spleens of WT^KL^ and C4^KL^ mice at the indicated ages. For **A-C**, each dot represents an individual mouse and bars represent means ± SEM. One-way ANOVA with Tukey’s post-test were performed and exact p values shown. For **(D**) multiple unpaired t-tests with false discovery rate (FDR) with two-stage step-up method (Benjamini, Krieger, and Yekutieli) was performed and exact p values are shown. For weeks 5-6, 7-10, 11-14, and 15-20: WT^KL^ n=17, 12, 4, 7 and C4^KL^ n=26, 10, 3, 9 respectively.

Given that both thymic output and homeostatic survival were increased in the C4^KL^ mice, we asked if homeostasis was completely dysregulated in the C4^KL^ mice, or if their CD4^+^ T-cell populations would plateau over time. Accordingly, we enumerated the total number of CD4^+^ T-cells in the lymph nodes and spleens of the C4^KL^ and WT^KL^ mice at 5-6, 7-10, 11-14, and 15-20 weeks of age. We found that the C4^KL^ mice had higher CD4^+^ T-cell numbers at each time point compared with the WT^KL^ mice, and peaked later, but that the numbers ultimately plateaued (**Figure 4D**). These data indicate that increased thymic output and homeostatic survival caused by the C4 mutation lead to partial dysregulation of peripheral CD4^+^ T-cell homeostasis.

### A higher frequency of CD4^+^ T-cells bearing the C4 mutant TCR respond to cognate pMHCII

We next asked how the C4 mutation impacts CD4^+^ T-cells responses to agonist pMHCII by monitoring their proliferation in response to immunization with cognate peptide antigen. Specifically, we adoptively transferred Tag-It Violet labeled WT^KL^, C4^KL^, or WT^CL^ CD4^+^ T-cells into C57BL/6J recipients, immunized the recipient mice with ovalbumin 323-339 (OVA) in Complete Freund’s Adjuvant (CFA), and quantified proliferation by Tag-It Violet dye dilution (**Figure 5A**). Enumeration of the daughter cells allowed us to back-calculate the number of responder cells as well as their proliferative capacity, meaning the average number of daughter cells that result from each responder(22). Our data indicate that the C4 mutation resulted in an increased number of CD4^+^ T-cells that responded to the cognate pMHCII by proliferating upon immunization but did not impact the proliferative capacity of those responders compared to the WT^KL^ and WT^CL^ CD4^+^ T-cells.

**Figure 5.**
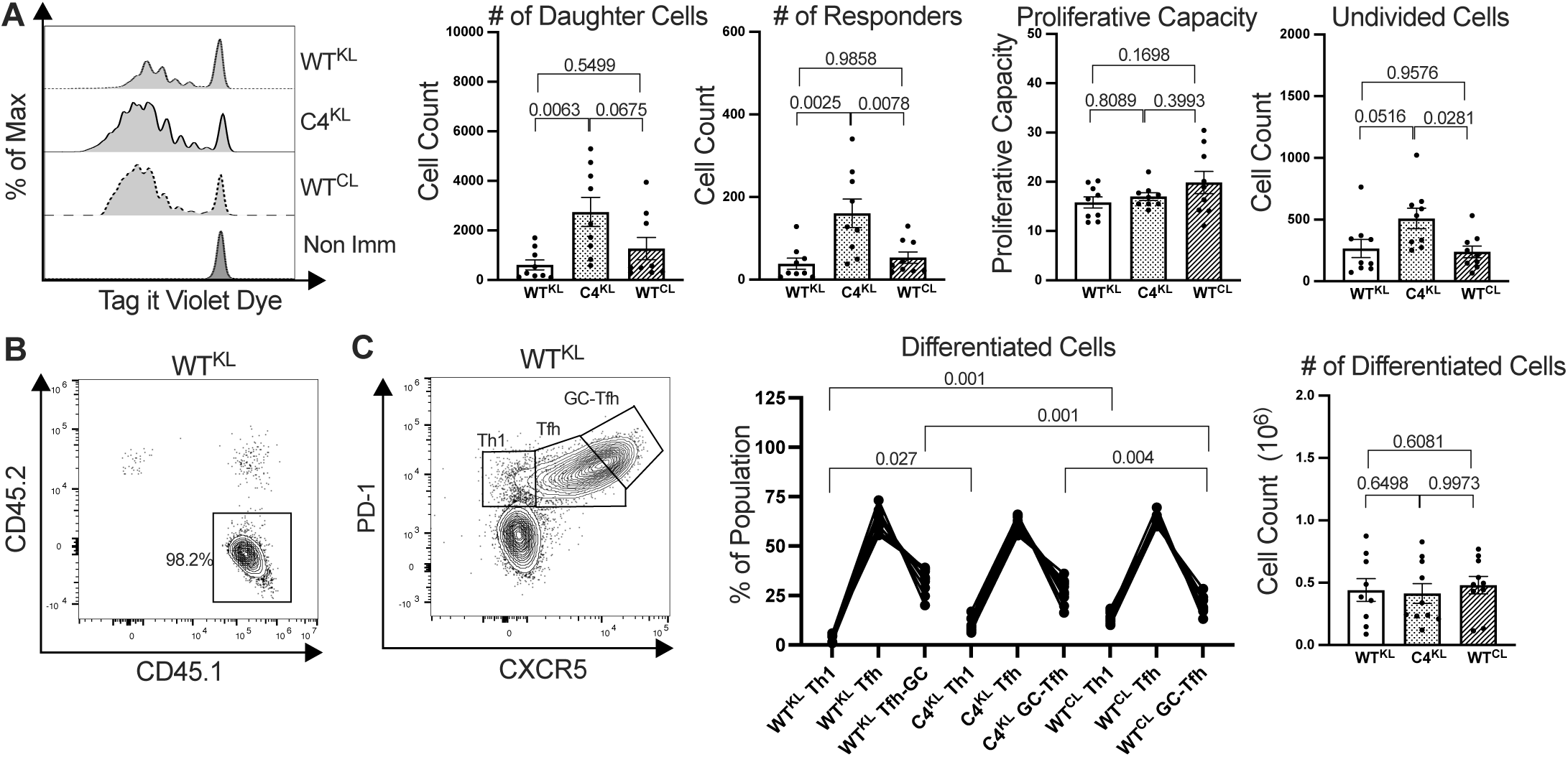
The C4 mutation does not impair CD4^+^ T-cell proliferation or differentiation. (**A**) (Left to right) Representative proliferation of WT^KL^, C4^KL^, and WT^CL^ CD4^+^ T-cells in C57Bl/6 recipient mice after immunization with OVA peptide in CFA, as measured by Tag-It Violet dye dilution. The number of divided cells, the number of responders calculated to generate the daughter cell population (see methods), the average number of daughter cells per responder (proliferative capacity, see methods), and the number of undivided cells are shown. Tag-It Violet cells were identified after FSC vs SSC, FSC-H vs FSC-A doublet discriminator, Live/Dead, CD4+CD8-, Vα2+Vβ5+, and CD45.1 (only for WT^KL^ and C4^KL^) or CD45.2 for WT^CL^. Each dot represents a recipient mouse (n=9). Bars equal means ± SEM. One-way ANOVA with Tukey’s post-test was performed and exact p values are shown. (**B**) Representative CD45.1 vs CD45.2 expression on WT CD4^+^ T-cells 6 days after immunization of recipient C57Bl/6 mice showing that the majority of cells after FSC vs SSC, FSC-H vs FSC-A doublet discriminator, Live/Dead, CD4^+^CD8^−^, Vα2^+^Vβ5^+^, CD44^hi^ gating are derived from the CD45.1^+^ adoptively transferred population. (**C**) Representative gating of adoptively transferred WT, C4, and Tac CD4^+^ T-cells 6 days after immunization of C57Bl/6 recipient mice for Th1 (CXCR5^lo^ PD-1^lo^), Tfh (CXCR5^Int^ PD-1^Int^), and GC-Tfh (CXCR5^hi^ PD-1^hi^) populations after pre-gating on FSC vs SSC, FSC-H vs FSC-A doublet discriminator, Live/Dead, CD4^+^CD8^−^, Vα2^+^Vβ5^+^ and CD44^hi^ (left). The percent of adoptively transferred populations differentiated to Th1, Tfh, and GC-Tfh are shown (middle) and the absolute number of adoptively transferred cells enumerated on day 6 are shown (right).

Because TCR signal strength has been shown to influence CD4^+^ T-cell differentiation, we next asked if the C4 mutation influences CD4^+^ T-cell differentiation on day 6 post-immunization as per prior work(23). To identify the transferred WT^KL^ and C4^KL^ CD4^+^ T-cells, we took advantage of their clonotypic TCR Va2 and Vb5 expression as well as the congenic CD45.1 marker. Given that >97% of the Va2^+^ Vb5^+^ CD44^hi^ CD4^+^ T-cells in recipient mice that received WT^KL^ CD4^+^ T-cells derived from the adoptively transferred population, we also used Va2^+^ Vb5^+^ CD44^hi^ gating to identify the WT^CL^ CD4^+^ T-cells since they did not express CD45.1 (**Figure 5B**). For all strains, we then used CXCR5 vs. PD-1 expression to identify Th1, T follicular helper (Tfh), and germinal center-Tfh (GC-Tfh) CD4^+^ T-cells. As previously reported, the WT^KL^ and WT^CL^ OT-II TCRs directed differentiation that heavily favored the Tfh phenotype, and this was also true for CD4^+^ T-cells bearing the C4 TCR(23). Small differences were observed between the frequencies of effector populations, but we could not confidently attribute these differences to the C4 mutation over strain-specific differences such as TCR levels. Finally, it is notable that by day 6 post-immunization, the number of responders was equivalent for the WT^KL^, C4^KL^, and WT^CL^ CD4^+^ T-cells, indicating that the differences in proliferation observed at 60hrs did not translate into more effector cells at day 6. These data indicate that the C4 mutation did not dramatically influence the responses of monoclonal populations of CD4^+^ T-cells against their cognate ligand.

## Discussion

The tertiary structure of αβTCR Cα domain, composed of the C- and F-strands, is conserved in mouse and man but deviates from the C-type Ig domains found in antibodies or γδTCRs, suggesting that it confers an evolutionary fitness advantage with regards to the function of CD4^+^ and CD8^+^ T-cells(24). Here we evaluate the fitness cost of acquiring mutations in this region for CD4^+^ T-cell development and homeostasis. Based on the phenotype of our mutant mice, we infer that the WT Cα surface of αβTCRs plays a role in regulating CD4^+^ T-cell development and homeostasis in response to self-pMHCII.

The data presented here contribute to our understanding of the features that shape the αβT-cell repertoire by ensuring that mature CD4^+^ T-cells are capable of distinguishing self-from foreign-peptide antigens presented in MHCII. They indicate that, for a monoclonal population of thymocytes expressing the MHCII-restricted OT-II TCR, mutating the C-strand results in enhanced positive selection without impacting negative selection, suggesting that the mutation increases sensitivity to weak TCR-pMHCII interactions. This conclusion is supported by phenotypic evidence that mature CD4^+^ T-cells expressing the mutant TCR experience higher tonic signaling in response to self-pMHCII and data showing they have increased homeostatic survival. We therefore infer from the phenotype of our mutant retrogenic and transgenic mice that a key function of the unique Cα domain surface is to regulate sensitivity to weak interactions with self-pMHCII. It is likely that the Cα domain similarly regulates CD8^+^ T-cell development and homeostasis, though confirming this will require future studies. Likewise, elucidating how the Cα domain impacts the composition and size of polyclonal CD4^+^ and CD8^+^ T-cell repertoires will require analysis with more sophisticated mouse models. Interestingly, this mutation did not dramatically impair responses to cognate antigen here or in our prior work; it is therefore possible that the unusual Cα surface evolved specifically to regulate responses to weak TCR interactions with pMHCII.

One caveat that should be mentioned here is that we observed distinct expression levels for the transgenic WT^KL^, C4^KL^, and WT^CL^ TCRs we evaluated. Both WT^KL^ and WT^CL^ CD4^+^ T-cells performed similarly in the periphery, despite having distinct TCR expression; therefore, given that the C4 TCR expression was between the WT^KL^ and WT^CL^ expression, differences in TCR expression are unlikely to account for the phenotypic differences observed in the periphery. We cannot fully rule out that the overlapping but distinct TCR expression between the WT^KL^ and C4^KL^ thymocytes influenced their phenotypes. Nevertheless, since two distinct pMHCII-restricted TCRs bearing the C4 mutation yielded phenotypes with an increased SP:DP ratio when compared to their counterpart in two different mouse model systems, the simplest interpretation of our data is that the thymocyte phenotype is largely mutation-intrinsic.

Prior work indicated that the unusual Cα surface is available to mediate protein-protein interactions, including homotypic TCR interactions that are impaired by the C-strand mutation of the Cα surface studied here(6-8). It is therefore notable that the F-strand is flanked by putative N-linked glycosylation sites in mouse and man while the human αβTCR has an additional glycosylation site in the middle of the C-strand(24). These sites might always be glycosylated, which could sterically hinder the proposed protein-interactions(25). Yet, for other proteins, varying glycosylation has been reported to regulate the activity of specific function-mediating surfaces. For example, antibody glycosylation in the Fc region is reported to alter their affinity for Fc receptors and thus serve as a mechanism to tune biological activity(26). It is therefore worth considering that differential glycosylation of the Cα surface may allow for regulation of TCR dimerization, or interactions with other proteins, to fine-tune signaling in response to weak TCR interactions with pMHCII. While future work will be required to explore this idea, the data presented here suggest that such regulation could aid in controlling thymic output and homeostatic survival of clonotypes in the periphery to regulate the size and diversity of the αβT-cell repertoire.

In closing, this study contributes to our understanding of CD4^+^ T-cell biology by providing insights into the relationship between a uniquely evolved structural feature of the TCR, the Cα domain, and the ability of CD4^+^ T-cells to distinguish self-from foreign-pMHCII and respond accordingly. While early debates about the nature of TCR triggering considered if TCR-CD3 clustering or conformational changes in the TCR-CD3 complex led to signal initiation, work in recent years has shown that the assembly of TCR-CD3-CD4/8 receptor complexes around pMHCI/II generates signals in response to single agonist pMHCI/II and involve conformational changes(1, 27-32). The data presented here help answer how the unusual Cα region contributes to TCR-directed responses by providing evidence that it has evolved to help the αβTCR mediate the unique function of responding to weak self-pMHCII by setting a threshold below which TCR-pMHCII interactions do not mediate positive selection or homeostatic survival.

## Materials and Methods

### Cell lines and constructs

M12 cell lines used as APCs for proximal signaling experiments that express tethered OVA:I-A^b^ or E641:IA^b^ have been previously described(33). OT-IIα cDNA were amplified by RT-PCR from OT-II transgenic mice and cloned into pUC18 using 5’ primers targeting the Vα2 leader sequences and 3’ primers targeting the C-terminal end, including the stop codon, of the TCRα and TCRβ constant regions, as previously described (19, 33). OT-IIβ was similarly amplified by RT-PCR from cDNA. Here we used a 5’ primer that targeted the Vb5 sequence immediately after the leader sequence as well as a 3’ primer targeting the C-terminal end of the Cβ2 region including the stop codon. The PCR product was cloned in-frame with sequence encoding the Vβ3 leader sequence in pUC18. The C4 (N170D and K173E, uniprot convention) mutations were introduced into the Cα domain by shuttling the 2B4 TCR Cα domain from the 2B4 TCR containing these mutations into the OT-II TCR by restriction enzyme cloning. After sequence verification in pUC18, the cDNA were subcloned into the hCD2 minigene cassette using standard techniques (34).

### Retrovirus Preparation

Phoenix E cells were cultured in DMEM 10%FCS supplemented with L-Glu and Pen-Strep. Cells were transfected using FuGene-6 with previously reported WT 2B4αTCR constructs or the C4 (N151D and K154E), F4 (F189E and K196A), or DE1 (K171A and D174A) mutants(6). Virus were harvested for two days and concentrated using 100kDa Amicon Ultra spin concentrator. Concentrated supernatants were used for retroviral transduction.

### Mice

To generate retrogenic mice we used 2B4βTg mice crossed to B10.A.*Rag2*^*-/-*^ mice as bone marrow donors (6). Mice were treated with 2mg 5-fluorouracil (5-FU) i.v. and bone marrow was harvested 5 days later. Hemopoietic stem cells were cultured overnight in IMDM, 5% fetal calf serum, pen/strep, L-Glu, NEAA, Pyruvate, 50mM 2-ME containing SCF, Flt3-L, and IL-11 cytokines 10ng/mL. Cells were spin infected with concentrated 2B4α WT, C4, F4, or DE1 loop mutant retrovirus in cytokine deficient media with 4mg/mL polybrene at 32°C for 1.5 hours.

Cells were washed and cultured overnight, then injected into sub-lethally irradiated (450rads) B10.A Rag2KO recipient mice at 6 weeks of age. T-cells were checked by tail bleeds 4 weeks post transfer. Thymus and spleen were analyzed 8 weeks post transfer on a Coulter flow cytometer. Mice were housed at Stanford University and used with approval of the Stanford University Committee on Animal Welfare. Note that retrogenic mice were made and analyzed in 2004-2005 and collected as .lmd files that cannot be read by current versions of FlowJo. For this reason, we have but percentages from previously analyzed data but not included flow plots. The data is intended to be preliminary and supportive of generating transgenic mice.

For the generation of OT-II TCR transgenic mice, the hCD2-OT-IIα WT or C4 constructs were co-injected with hCD2-OT-IIβ constructs into C57BL/6 embryos at the University of Arizona’s Genetically Engineered Mouse Model Core (GEMMCore) facility. Founder mice were crossed to C57Bl/6J Rag1KO mice (Jax, Strain Number 002216 B6.129S7-Rag1^tm1Mom^/J), and then crossed to C57Bl/6J Rag1KO CD45.1 congenic mice (a generous gift of the Nikolich-Zugich lab) to generate OT-II WT or C4 C57Bl/6J CD45.1 Rag1KO mice.

These mice were then backcrossed with C57Bl/6J Rag1KO CD45.1 mice to generate hemizygous mice for analysis. C57Bl/6J mice (JAX Strain:000664) used as recipients in adoptive transfer experiments were purchased from the Jackson Laboratory, Bar Harbor, Maine. OT-II TCR transgenic mice and recipient mice were maintained under specific pathogen-free conditions in the animal facility at the University of Arizona. Experiments were conducted under guidelines and approval of the Institutional Animal Care and Use Committee of the University of Arizona.

### Adoptive transfers, peptides, CFA, and immunizations

From male mice, brachial, axillary, and inguinal lymph nodes were pooled with spleens and CD4^+^ T-cells were enriched using a CD4 T-cell isolation kit (Miltinyi Biotec) and MACS separation columns (Miltinyi Biotec). 1×10^5^ Tag-it Violet (BioLegend) labeled CD4^+^ T-cells were retro-orbitally injected into male C57BL/6J mice recipient mice. 5µM Tag-it Violet staining was performed according to manufacturer’s instructions. One day later, mice were immunized with 20µg ovalbumin (OVA) peptide (323-339) (ISQAVHAAHAEINEAGR, purchased at >95% purity from 21^st^ Century Biochemicals, Marlborough, MA.) in Complete Freunds Adjuvant (CFA, Sigma-Aldrich, St. Louis, MO.) on the back flank. At 60 hours (proliferation analysis) or 6 days (differentiation analysis) post-immunization the draining lymph nodes (dLNs: brachial, axillary, inguinal) were harvested and CD4 enriched prior to antibody staining and flow cytometry analysis.

### Proliferation Analysis

T-cells labeled with Tag-It Violet and subsequent dilution of fluorescent dye detected by flow cytometry were used to calculate the total number of daughter and undivided cells, the number of responding T-cells (number of original T-cells that divided due to stimulus), and the proliferative capacity (the average number of daughter cells generated per responder) as described similarly(22).

### Flow Cytometry

Single-cell suspensions were blocked with anti-mouse FcRII mAb clone 2.4G2 hybridoma supernatants (ATCC), and then stained for 30 min at 4°C with corresponding Abs. Staining thymocytes or spleenocytes included anti CD4 (RM4-5), CD8α (53-6.7), CCR7/CD197 (4B12), TCR Vα2 (B20.1),TCR Vβ5.1/5.2 (MR9-4), CD69(H1.2F3), PD-1 (29F.1A12), CD5 (53-7.3),Ly6C (HK1.4), CD8β (YT5156.7.7), CD62L (MEL-14), Qa-2 (695H1-9-9), CD24 (M1/69), CD25 (PC61), CD44 (IM7), CD45.1 (A20), CD45.2 (104), CD185/CXCR5 (L138D7) all purchased from BioLegend. Staining for CCR7/CD197 was performed at 37°C, 30 mins prior to surface stain.

Cells were then stained using LIVE/DEAD Blue (Invitrogen) 30 min at 4°. Cells were fixed using 4% paraformaldehyde for 15 minutes at 4°C. Cells were washed before being stored at 4°C. If applicable cells were permeabilized the following day using BD Bioscience Perm/Wash buffer for 30 min at 4°C. Cells were washed with Perm/Wash buffer twice before stained with rabbit anti-Cleaved Caspase 3 (Asp175) (Cell Signaling Technologies) at a 1:200 dilution for 30 mins at room temperature and was detected with anti-rabbit BV421 (Dnk, BioLegend). Flow was performed on Cytek Aurora (Cytek) and analyzed on Flowjo V10 (Becton, Dickinson & Company).

### Intracellular signaling analysis

Basal phosphorylation ζζ module of the TCR-CD3 complex was measured by intracellular staining. Naïve WT^KL^ C4^KL^ or WT^CL^ inguinal lymph nodes were harvested, immediately processed into single cell suspensions, and fixed with 4% paraformaldehyde for 15 minutes at 37°C. The lymphocytes were then blocked with anti-mouse FcRII mAb clone 2.4G2 hybridoma supernatants (ATCC) for 30 minutes on ice. Cells were stained with anti-CD4 (GK1.5, Biolegend) and fixed with 4% paraformaldehyde for 15 minutes at room temperature. Cells were washed twice with FACs buffer, pelleted at 350xg for 5 minutes at room temperature, resuspended in 1mL True-Phos Perm Buffer (BioLegend Inc), and incubated at -20°C for 2 hours. They were then stained for 60 minutes with anti-pCD3ζ (K25-407.69, BD Biosciences) at 37°C, washed, and analyzed on a LSR II (BD Biosciences) at the Flow Cytometry Shared Resource at the University of Arizona.

To measure pζζ levels upon stimuli, enriched naïve WT^KL^ CD4^+^ T-cells were labeled with Tag-It Violet (BioLegend Inc) while E641:I-A^b+^ (null) or Ova:I-A^b+^ (cognate) M12 cells expressing tethered pMHCII complexes were labeled with CFSE (ThermoFisher) according to the manufacturer’s instructions. M12 cells and naive CD4^+^ T-cells were chilled on ice for 30 minutes, 5×10^5^ of each cell type were mixed in 1.5ml snap cap tubes, and the cells were pelleted at 2000 rpm for 30 second at 4°C to force interactions. The supernatant was removed and the tubes were transferred to a 37°C water bath for 2 minutes to enable signaling. Fixation Buffer (BioLegend Inc) was then added for 15 minutes at 37°C. Cells were washed twice with FACs buffer, pelleted at 350xg for 5 minutes at room temperature, resuspended in 1mL True-Phos Perm Buffer (BioLegend Inc), and incubated at -20°C for 2 hours.

Cells were blocked with anti-mouse FcRII mAb clone 2.4G2 hybridoma supernatants (ATCC) for 30 minutes, pelleted, and stained on ice for 60 minutes with anti-pCD3ζ (K25-407.69, BD Biosciences). Finally, cells were washed 2x with FACs buffer at 1000xg for 5 minutes at room temperature and analyzed on a LSR II (BD Biosciences) at the Flow Cytometry Shared Resource at the University of Arizona. 5×10^4^ CD4^+^ T-cell:M12 cell couples were collected per sample. Flow cytometry data were analyzed with FlowJo Verison 10 software (Becton, Dickinson, & Company) by gating on T-cell and M12 cell couples, as described previously (10).

## Acknowledgements

We thank Mark M. Davis for providing critical feedback and use of the retrogenic mouse data. We thank J. Nikolich-Žugich for providing Rag1KO CD45.1 C57BL/6J mice and Eric Huseby for providing the hCD2 cassette. We also thank Dominik Schenten, Koenraad Van Doorslaer, and members of the Kuhns Lab for critical feedback on the manuscript.

## Supplemental Figure Legends

**SFigure 1:**
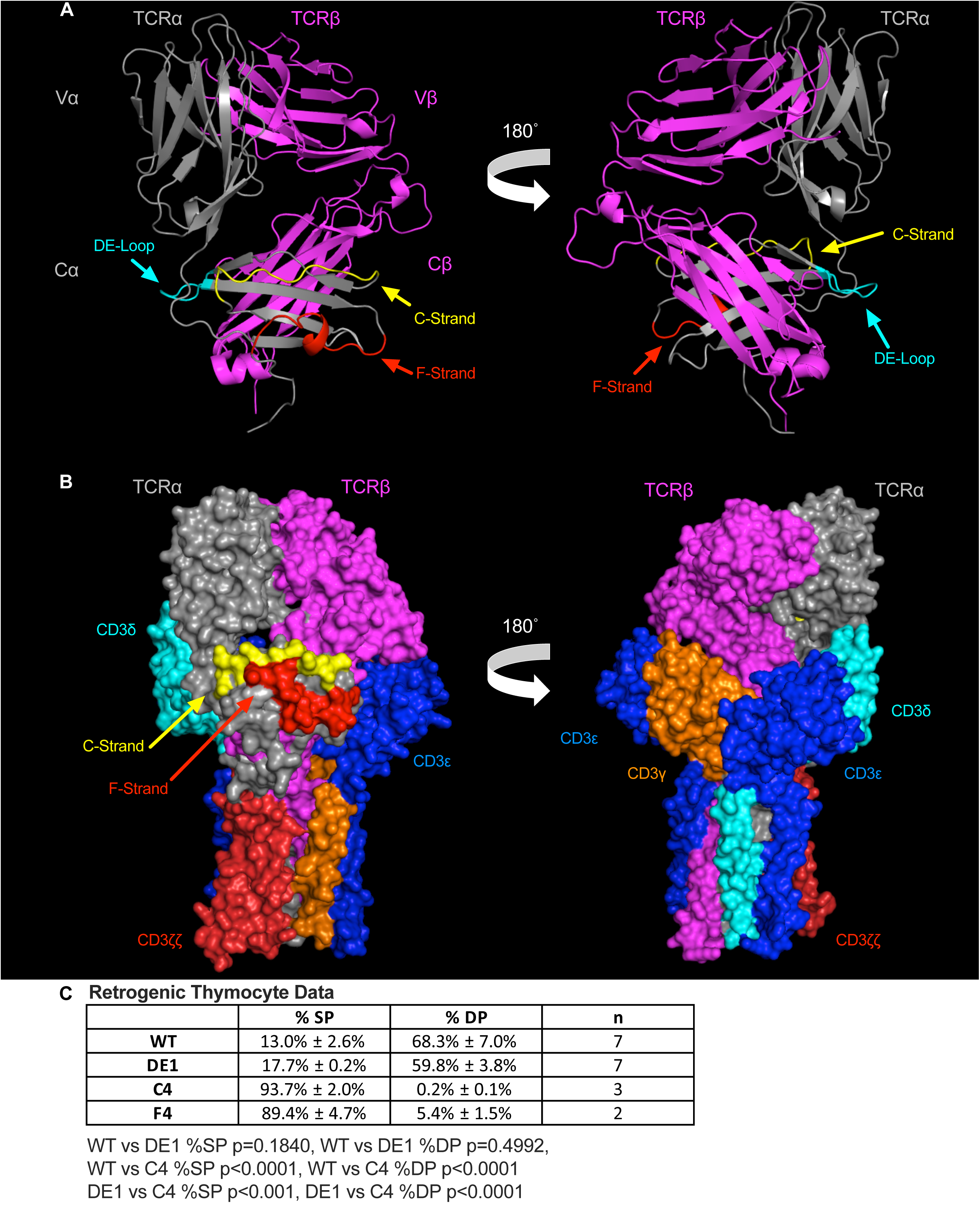
Functional sidedness of the αβTCR. (**A**) Cartoon representations of the αβTCR showing the side of the TCR with the unusual Cα surface (left) and the side where CD3γε and CD3δε dock (right)(PDB: 1TCR). (**B**) Cryogenic electron microscopy (cryo-EM) spacefill representation of αβTCR-CD3 complex showing the free side of the αβTCR with the exposed Cα surface (left) and the side with docked CD3 modules (right) (PDB: 6JXR). For both (**A**) and (**B**), structural features are colored as followed: TCRα (grey), TCRβ (magenta). The C-strand (yellow), F-strand (red) and DE-loop (cyan) are highlighted on the TCRα constant region (Cα). CD3 modules are colored as CD3δ (cyan), CD3ζζ (red), CD3ε (blue), and CD3γ (orange) are highlighted. (**C**) DP (CD8^+^ CD4^+^) and SP (CD8^−^ CD4^+^) thymocyte percentages are shown for retrogenic mice with the mutated 2B4 TCR. Values are shown as percentages ± SEM, n is number of mice per group. Statistical analysis was done using One-way ANOVA and Tukey’s post-test with exact p values shown. F4 was not included in the analysis due to insufficient n.

