## Supplemental Figure 1 for "The TCR Cα domain regulates responses to self-pMHCII"

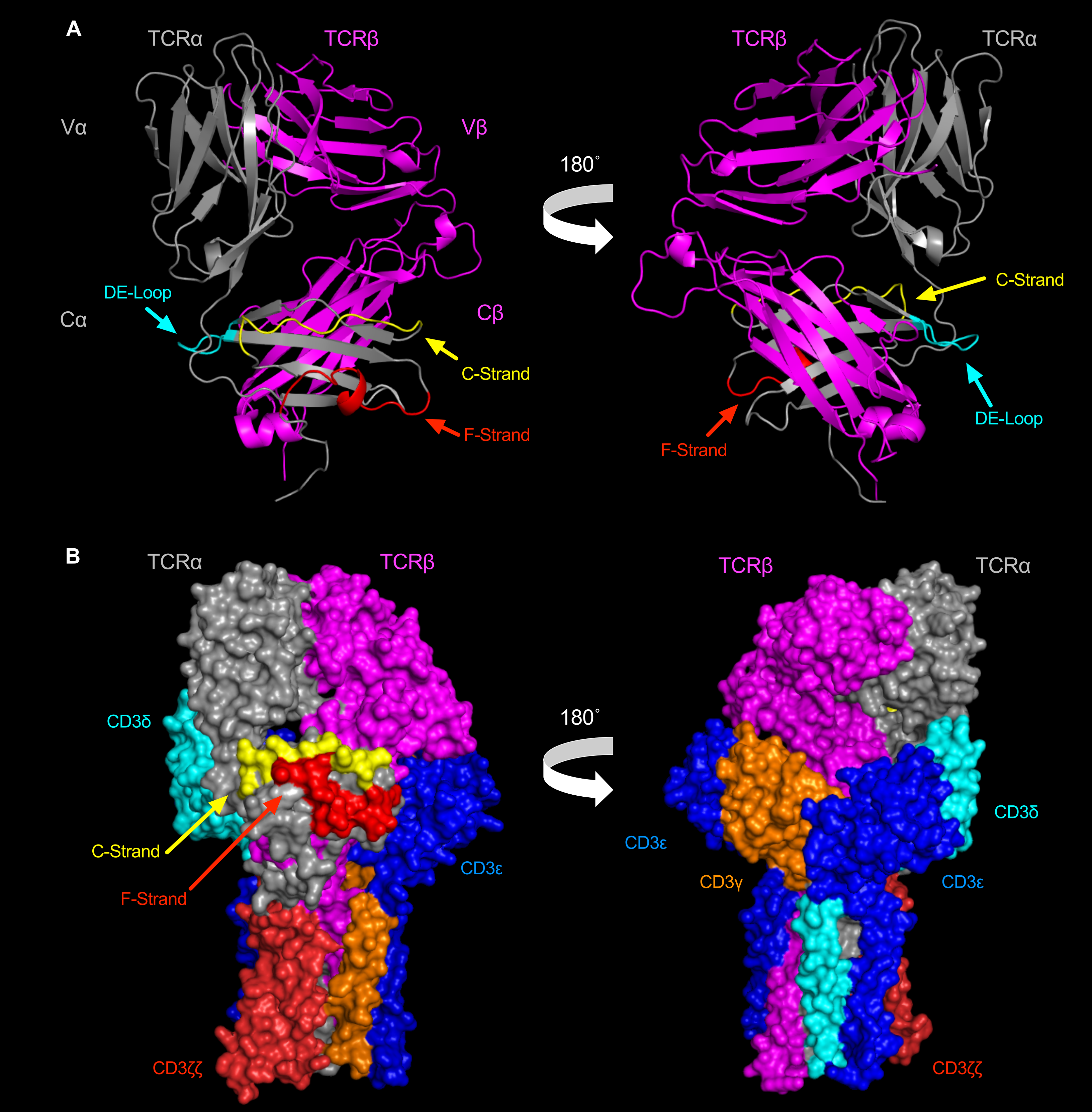


**C** Retrogenic Thymocyte Data

| | % SP | % DP | n |
|---|---|---|---|
| **WT** | 13.0% ± 2.6% | 68.3% ± 7.0% | 7 |
| **DE1** | 17.7% ± 0.2% | 59.8% ± 3.8% | 7 |
| **C4** | 93.7% ± 2.0% | 0.2% ± 0.1% | 3 |
| **F4** | 89.4% ± 4.7% | 5.4% ± 1.5% | 2 |

WT vs DE1 %SP p=0.1481, WT vs DE1 %DP p=0.4401,
WT vs C4 %SP p<0.0001, WT vs C4 %DP p<0.0001

**Supplemental Figure 1**
